# Taxonomic survey of *Anadenobolus monilicornis* gut microbiota via shotgun nanopore sequencing

**DOI:** 10.1101/560755

**Authors:** Orlando J. Geli-Cruz, Matias J. Cafaro, Carlos J. Santos-Flores, Alex J. Ropelewski, Alex R. Van Dam

## Abstract

Millipedes constitute one of many soil-inhabiting organisms that act as important components of litter decomposition and nutrient recycling in terrestrial ecosystems. This is thanks in part to the microbial diversity that they contain in their gut compartments. However, millipedes and their gut microbiota are understudied, compared to other arthropods. For this reason, we partook in a metagenomic analysis of the gut of *Anadenobolus monilicornis*. We collected specimens of *A. monilicornis*, which were starved for a varying amount of time, from different municipalities of Puerto Rico. Once the DNA from their guts was extracted and sequenced using the MinION nanopore sequencer, we proceeded to analyze and compile the data obtained from the sequencer using programs such as Phylosift and MEGAN6 and the web-based MG-RAST. From our two best samples, we obtained a total of 87,110 and 99,749 reads, respectively. After comparing the data analyses and gene annotation done for both samples, we found that the bacterial phyla Proteobacteria, Bacteroidetes and Firmicutes were consistently well represented; one of our samples had much more Chlamydiae representation than the other, however. Sampled eukaryote phyla include Arthropoda, Chordata and Streptophyta. We would need a greater sample size to better determine differences in microbial diversity between millipede populations across the island; considering our small sample size, however, we were able to broadly reveal the diversity within the microenvironment of *A. monilicornis*’s gut.

## Introduction

Millipedes are a group of arthropods belonging to the class Diplopoda and the subphylum Myriapoda. With around 12,000 described species worldwide, they are a diverse group found in a variety of habitats from humid rainforests to xeric deserts [1–3]. Millipedes are one of many soil-inhabiting decomposers, alongside other macroinvertebrates such as earthworms and isopods [4–6]. They are considered in some ecosystems as one of the more important components of terrestrial litter decomposition and nutrient cycling [4–6]. However, they are an understudied arthropod group and have received little attention to elucidate their interactions with their gut microbial community, despite their significance in nutrient cycling at the soil leaf litter interface [7,8].

In comparison, the microbial composition of the gut of leaf litter macroinvertebrates has received some attention from the scientific community. Termites, for example, are known to harbor diverse clades of bacteria from the phyla Bacteroidetes, Firmicutes and Spirochaetes in their guts, depending on the diet and gut compartments studied [9]. The guts of terrestrial isopods, which also have a similar ecological role to millipedes, are well represented by Proteobacteria [10]. In a similar fashion, millipedes have been shown to host certain microorganisms in their guts which aid in digestive processes [11]. Dilution plating techniques have shown that some millipede species harbor an abundant diversity of proteobacteria and actinobacteria [12].

There have been a small number of microbial community surveys of the gut of millipedes. In some species, the most dominant bacteria were found to belong to the Enterobacteriaceae family; in addition, ascomycetes were the most common yeast strains found [13]. A desert-dwelling millipede species possesses gut bacteria that can degrade cellulose, contributing to nutrient cycling in deserts [1]. As well, certain species from the millipede orders Julida, Spirobolida, and Spirostreptida harbor an association between methanogenic archaea and ciliate protozoa in their hindguts, contributing to methane production [14]. The diversity of bacteria and other microorganisms that occur in millipede guts might be of interest to field ecologists and microbiologists alike, as the interactions between these organisms affects both soil nutrient recycling and organic matter decomposition [15,16]. A full genetic or metagenomic approach to these problems, however, has yet to take off.

For this study, we will be focusing on the microbiota that inhabits *Anadenobolus monilicornis*’s digestive tract. *A. monilicornis* is a species native to the Caribbean which has also been introduced to Florida (U.S.A.), where it is treated as a pest [17,18]. It is considered the most common millipede in the karst zones of Puerto Rico [3]. Previous microbiota studies have been morphohology based for *A. monilicornis*; specifically, Contreras & Cafaro [19] conducted a morphometric study of the protozoa *Enterobryus luteovirgatus*, which forms a commensalistic relationship with the millipede. Beyond this study, very little is known about this species.

To identify the microbial diversity inside *A. monilicornis*’s gut, we will utilize a shotgun metagenomic analysis. Shotgun sequencing of environmental samples, most notably from microbial communities [20], can lead to the discovery of microorganisms that are otherwise difficult or impossible to culture in a laboratory setting [21]. Several metagenomic sequencing studies have successfully revealed complex host-symbiont relationships, such as those that occur in deep-sea tube worms [22], termites [23] and Daphnia species [21]. Particularly, metagenomic studies involving arthropods have focused mostly on insects [24–27], successfully revealing their microbial diversity. However, there are few comprehensive metagenomics sequencing surveys of the microbiota of other non-insect arthropods, millipedes included [28,29].

To analyze the DNA of *A. monilicornis*, Oxford Nanopore Sequencing Technology (ONT) will be used. This relatively new technique works by identifying the order of nucleotides in a DNA sequence as it passes through individual nanopore channels. ONT has already proven useful for producing relatively long DNA sequences, real-time analysis, and detection of structural sequence variants [30]. Specifically, we will be using the MinION, a portable nanopore sequencer developed by Oxford Nanopore Technologies. Our objective is to begin to develop protocols for shotgun sequencing of millipede guts via ONT, and to develop an initial fingerprint of the microbial diversity found within *A. monilicornis* using ONT as a tool to elucidate this complex system.

## Materials and Methods

### Millipede Sampling

*Anadenobolus monilicornis* millipedes were collected from two locations collected in July 2017 from western Puerto Rico: The University of Puerto Rico, Mayagüez Campus, and the municipality of Rincón. These were kept alive and starved for 24 hours (Mayagüez samples), and ten days (Rincón samples). The millipedes were collected and kept in small glass containers with moist filter paper. The Rincón sample was kept until it began to eat the filter paper and the frass showed little signs of plant material.

### Gut and DNA extraction

The gut extraction and DNA extraction work were done in the Symbiosis laboratory at the University of Puerto Rico, Mayagüez Campus. Following workstation and lab material sterilization with 10% bleach, the head and the last two or three segments of the abdomen of the specimens were cut and removed with a scalpel; the abdomen was cut to facilitate gut extraction. The guts were removed and placed in 2mL tissue disruption tubes, where they were liquified by manually shaking the tubes.

We followed the Qiagen Fast DNA Tissue Kit (cat. No. 51404) protocol to purify the DNA samples from the specimens. The buffer for the Qiagen Fast DNA Tissue Kit was prepared before use as the protocol specified: 40 mL of ethanol were added to the AW1 and AW2 Buffer concentrates, and 25mL of isopropanol were added to the Buffer MVL concentrate. The tubes were spinned down briefly via vortex mixer and set on a block heater at 56°C for one hour. The tubes were briefly spinned down every 10 minutes for that hour. 265µL of the Buffer MVL mixture (200µL of AVE, 40µL of VXL, 20µL of Proteinase K, 1µL of DX Reagent and 4µL of RNase A) were added to the tubes.

For all samples, the mixtures were moved to a QIAamp Mini Spin Column and centrifuged for one minute at 15,000 rpm. The spin column was then placed in a clean 2mL collection tube, while the previous tube and filtrate were discarded. 500µL of the AW1 Buffer were added to the spin column before being centrifuging, again for one minute at 15,000 rpm; the spin column was placed in another 2mL collection tube, and the previous tube discarded. 500µL AW2 Buffer were added to the spin column before centrifuging. The spin column was then added into a new collection tube, which was centrifuged again for two minutes, and later placed in a clean 1.5mL microcentrifuge tube. 50µL of nuclease-free water was added directly into the spin column, which was left subsequently for one minute at room temperature and later centrifuged for one minute. This last step was repeated once to increase yield. After this step, the Oxford Nanopore Technologies (ONT) 1D PCR barcoding genomic DNA (SQK-LSK108) for version R9 chemistry procedure was followed, with some minor alterations.

### DNA Fragmentation

A master mix of 14.14µL of Fragmentase buffer and 2.2µL of 10X NEBNext® dsDNA Fragmentase® (NEB cat. No. M0348s) was mixed first. In new tubes, we added 32µL of the samples and 8µL of the master mix to each. The new tubes were vortexed for two seconds and spun down; they were then placed on a thermocycler for five minutes at 37°C followed by approximately five minutes at 4°C. In order to heat kill the Fragmentase 5µL of EDTA was added and placed on a thermocycler for 15 minutes at 65°C followed by 10 minutes at 5°C. We aimed to produce 5,000-30,000Kb DNA fragments. DNA quality was checked using 2µL of each sample mixed with 3µL of loading dye and then added to a 1X gel set to 66V for 30 minutes.

Leftover enzymes were cleaned via Agencourt® Ampure® XP beads: 50µL of samples of each sample were added to 90µL of Ampure XP beads, mixed 10 times by pipetting. The mixture was left at room temperature for five minutes, then placed on a magnetic rack for two minutes. The cleared solution was then aspirated out. The process was repeated but with 200µL of 70% ethanol followed by aspiration. Finally, 48µL of nuclease-free water was added, and aspirated out into new 1.5 mL tubes and carried forward in the protocol.

### NEBNext FFPE DNA Repair

DNA was repaired via the NebNext FFPE DNA repair kit. 5.5µL of nuclease-free water, 6.5µL of FFPE DNA repair buffer, 2µL of NEBNext FFPE DNA repair mix (NEB cat No. M6630) and 53.5µL of the sample DNA were mixed. The samples were transferred to 0.2mL tubes for and placed in a thermocycler programmed to 20°C for 15 minutes, followed by 4°C for 10 minutes. The process ended with the previous Ampure XP beads cleaning procedure.

### NEBNext Ultra II End Repair / dA-Tailing module

We used the NEBNext® Ultra™ II End Repair/dA-Tailing Module (NEB cat No. E7546). 5µL of nuclease-free water, 3µL of NEBNext Ultra II End Prep Enzyme mix, 7µL of NEBNext Ultra II End Prep Reaction buffer and 45µL of sample DNA were placed in 0.2mL tubes. The tubes were transferred to a thermocycler programmed for 20°C for 20 minutes, followed by 65°C for 5 minutes and finally 4°C for a few minutes. The process ended with another round of Ampure XP bead cleanup as before.

### Ligation of Adapters

Ligation of Barcode Adapters was performed with NEB Blunt/TA Ligase Master Mix (NEB cat No. M0367). 20µL of ONT ligation adapter (ONT cat No. Ligation Sequencing Kit SQK-LSK108), 50µL of NEB Blunt/TA Ligase master mix and 30µL of sample DNA were mixed by inversion in a 1.5mL tube. The tube was left at room temperature for 10 minutes, followed by AMPure XP beads cleaning procedure The finished samples were then transferred to PCR tubes.

### Barcoding PCR

2µL of PCR barcode PCR Barcoding Kit (ONT cat No. SQK-PBK004), 2µL of 10ng/µL adapter ligated template, 50µL of NEB LongAmp Taq 2X Master Mix (NEB cat No. M0287), and 46µL of nuclease-free water. The samples were placed on a thermocycler using the following cycling conditions: 95°C for three minutes for initial denaturation, 95°C for 15s for denaturalization, 62°C for 15s for annealing and 4°C on hold. The process ended with the Ampure XP beads cleanup procedure as before.

### SpotOn Flow Cell Prep

We followed the ONT for the SpotOn Flow Cell version R9 chemistry (ONT cat No. FLO-MIN 107 R9): After extracting the buffer from inside the flow cell’s priming port by pipetting, we mixed 480µL of Running Buffer with Fuel (RBF) mix with 520µL of nuclease-free water and added 800µL of this mixture into the priming port via pipette. 35µL of RBF with 2.5µL of nuclease-free water, 25.5µL of Library Loading Bead kit (ONT cat No. EXP-LLB001) and 12µL of the DNA library were mixed. 200µL of the priming mixture (RBF & nuclease-free water) was loaded into the flow cell via the priming port by pipetting, while 75µL of the sample were loaded via the sample port in a dropwise fashion. The MinION was connected to a local MacBook, and the MinKNOW software program was accessed to start a sequencing run for 48 hours. Having obtained the data in the form of shotgun single long reads, we used the MinKNOW software to acquire and analyze the sequencing data obtained from the MinION. The libraries were then sequenced again on a second Flow Cell.

### Bioinformatic Analyses

#### Quality filtering and de-multiplexing

We used the MinKNOW software program for initial quality filtering of reads. The HDF5-formatted data from the Nanopore was moved from the MacBook to the Pittsburgh Supercomputing Center’s (PSC) Bridges Supercomputer. Within the PSC and using the Anaconda and Python environment, we installed the albacore basecaller v-2.1.3 [31], to separate the different barcodes and convert the data to FASTQ format [32]. Data can be found on NCBI SRA (BioProject# PRJNA521026). We used pauvre to investigate the overall quality and distribution lengths of our reads [33] (Fig 1). Next we used Trimmomatic as a second quality filtering step [34]. We used KmerGenie to predict k values for our datasets in order to attempt optimizing the genome assembly process [35]. We ran velvetg and velveth to attempt a *de novo* genome assembly [36]. This assembly was unsuccessful. We also tried Canu [37] but it also didn’t produce any scaffolds. The ONT data didn’t have sufficient depth of coverage to produce a *de novo* assembly. To summarize the diversity and relative abundance of the community of microbes sequenced we used a variety of metagenomic classification programs for our long-read data. We chose to use programs that should work well with shotgun long-read data produced by the ONT MinION sequencer.

**Fig 1.**
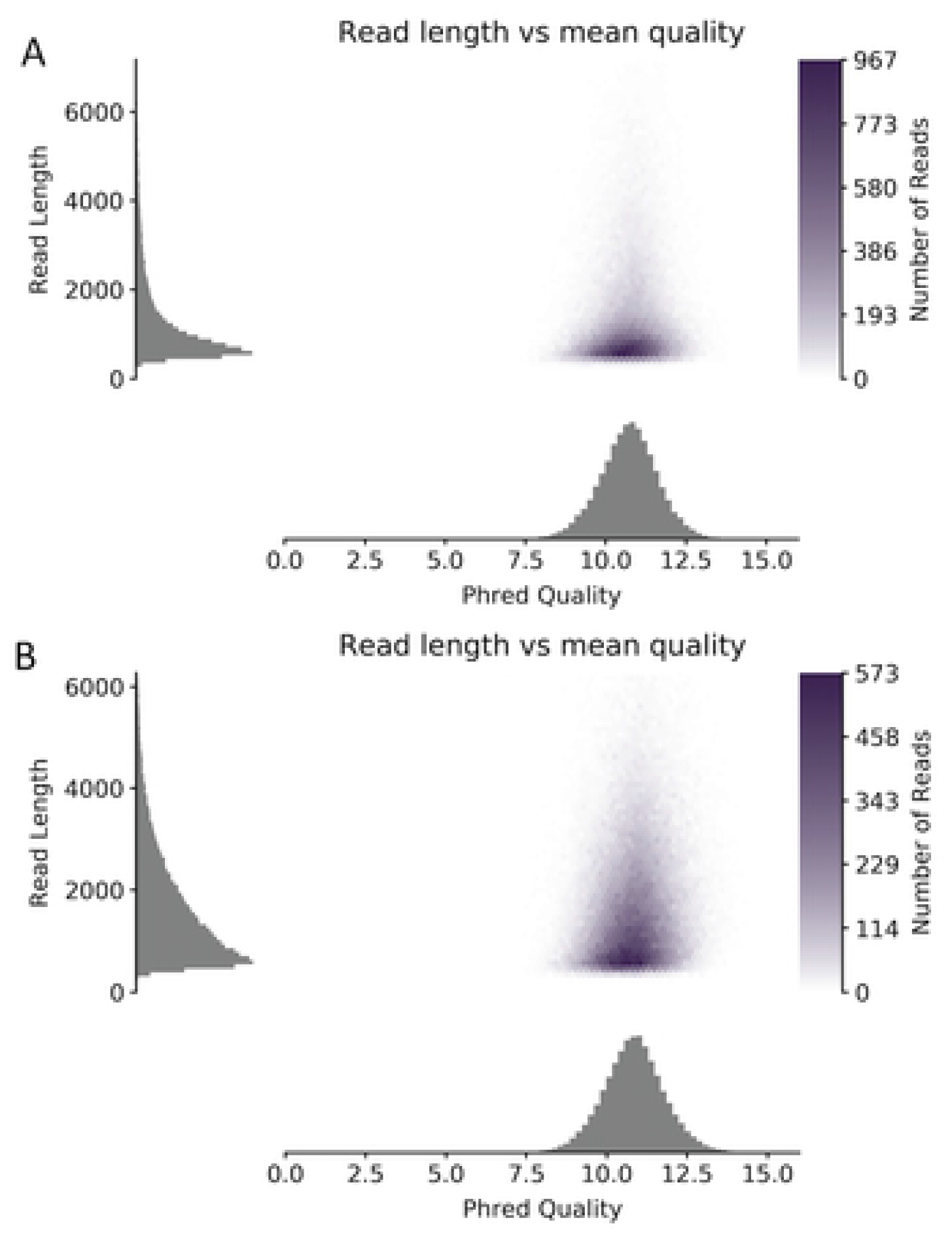
Read length and quality distribution of ONT nanopore reads produced on the MinION via R9 chemistry from *Anadenobolus monillicornis* gut samples. (A) Gut sample taken from Rincón and (B) gut sample taken from Mayagüez Puerto Rico.

#### Metagenomic Analyses

We used Phylosift to reconstruct a phylogenetic tree and place our organisms found in the sample [38]. We also used BLASTn to align the sequenced data to NCBI nt database [39]. We imported the data to MEGAN6 to do taxonomic, functional and comparative analysis of the data [40]. Finally, we uploaded our data to the MG-RAST server to analyze the metagenome and annotate the genes to their respective organisms and to metabolic processes; to compliment the latter, we also uploaded our sample data to GenomeNet obtain a KO lists. KO lists were then uploaded to iPath3 to analyze the sampled metabolic pathways [41,42].

## Results

### Quality filtered reads and summary statics

We were able to de-multiplex the two millipede gut samples via albacore, to which we will be referring to as the Mayagüez and Rincón samples. For the Mayagüez sample, we obtained a total of 87,110 quality-filtered reads; for the Rincón sample, a total of 99,749 reads. For more summary statistics, refer to Table 1. Phylosift matched 298 reads (261 bacterial, 36 eukaryotic) for the Mayagüez sample, and 48 reads (45 bacterial, 2 eukaryotic) for the Rincón sample. MG-RAST assigned taxonomic groups to 1,277 total reads for the Mayagüez sample, and 780 total reads for the Rincón sample. Finally, MEGAN6 was able to utilize 4,698 total reads, of which 3117 were assigned to taxonomic groups for the Mayagüez sample, and for the Rincón sample 5,626 reads were utilized with taxonomic groups assigned to 3,197 reads.

**Table 1.** Summary statistics for the quality-filtered *A. monilicornis* sample data.

|  | Mayagüez sample | Rincón sample |
| --- | --- | --- |
| Reads | 87,110 | 99,749 |
| Base Pairs | 132,196,067 | 176,209,113 |
| Mean Length | 1517.6 | 1766.5 |
| Median Length | 959.0 | 1388.0 |
| Min. Length | 187 | 137 |
| Max. Length | 18,142 | 14,296 |

The data overall was of phred quality scores were decent for nanopre data with the majority of the reads with a with a phred quality score greater than 9, i.e. 80% base call accuracy or better (Fig 1). However, the read distribution length was much shorter than expected and we had few reads that were approximately 10kbp in length (Fig 1). The results from this was that we were able to MEGAN6 produced a summary taxonomic tree of the phyla sampled showing the distribution of reads across phyla (Fig 2). Finally, the taxon accumulation curve created via MEGAN6 (Fig 3) starts to plateau around 20 phyla for the Mayaguez sample, and around 15 phyla for the Rincón sample, indicating that we will most likely discover 15 or more phyla in total with continued sampling.

**Fig 2.**
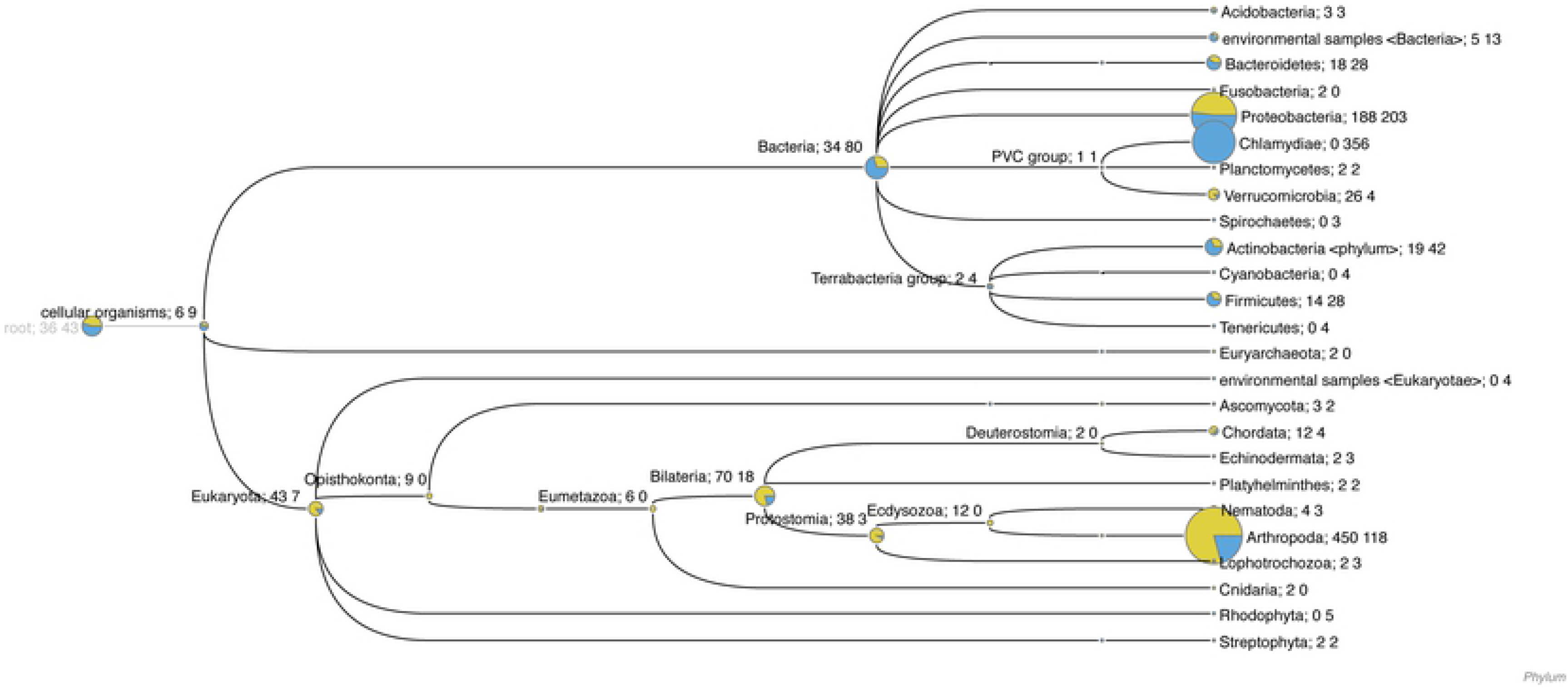
Sampled phyla representation from the Mayagüez sample (blue) and the Rincón sample (yellow), using MEGAN6.

**Fig 3.**
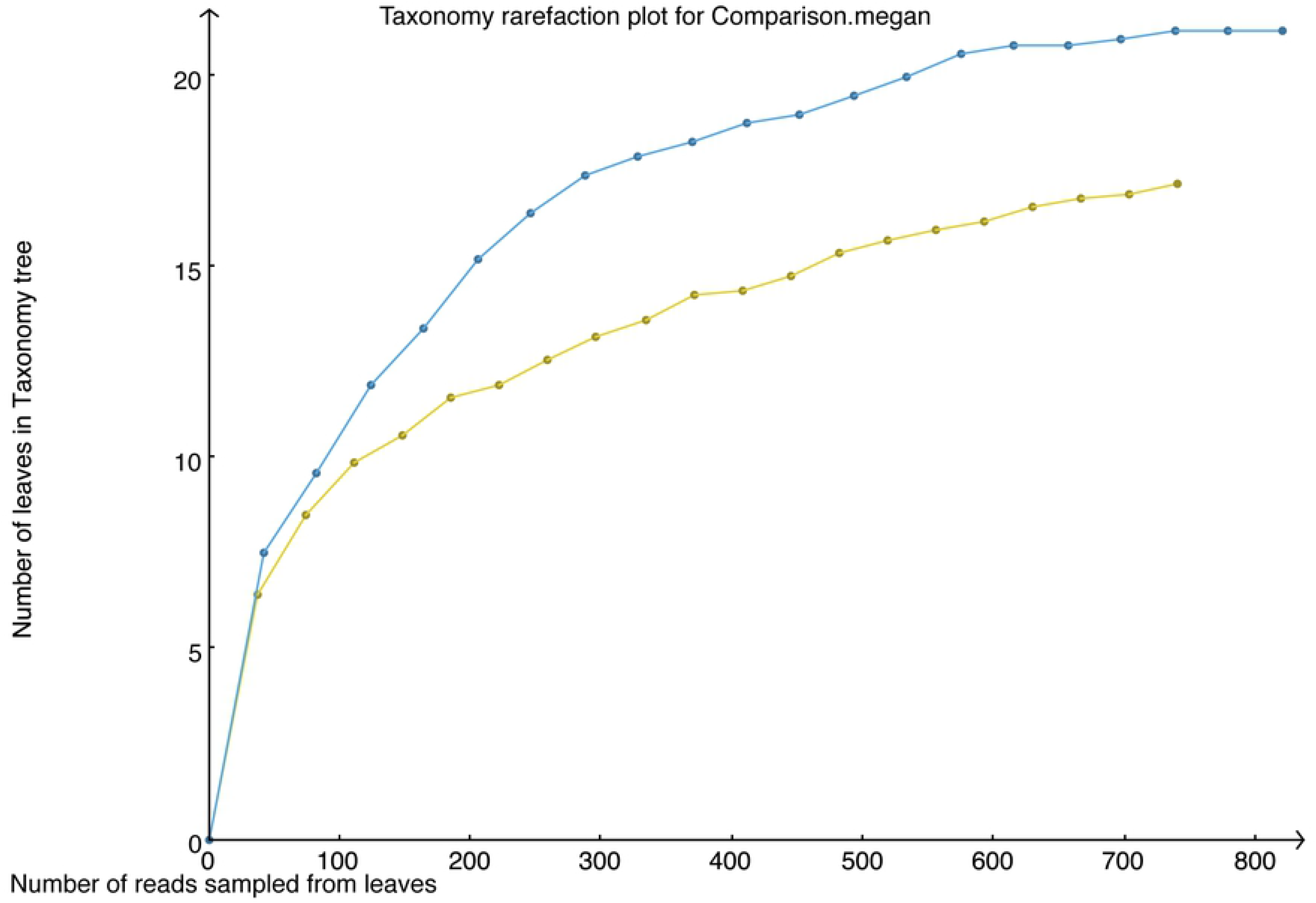
Taxonomic accumulation curve for the Mayagüez sample (blue) and the Rincón sample (yellow), created using MEGAN6.

### Bacterial Reads

Across the three metagenomics summary analyses (Phylosift, MEGAN6 and MG-RAST), the two samples showed similar bacterial phyla diversity: both samples had Proteobacteria, Firmicutes, and Bacteroidetes representation. According to the Phylosift, MG-RAST and MEGAN6 analyses, the Mayagüez sample had Chlamydiae as the most abundant phylum with a total of 187 reads for Phylosift (Fig 4), 673 reads for MG-RAST (Fig 5) and 356 reads for MEGAN6 (Table 3). The Rincón sample had Bacteroidetes and Proteobacteria as the most well represented phyla, with a total of 15 and 10 reads for Phylosift (Fig 4), 147 and 204 reads for MG-RAST (Fig 5), and 24 and 224 reads for MEGAN6 (Table 2), respectively. Phylosift indicated that Bacteria represented 87% of the sampled reads for the Mayagüez sample, and 96% for the Rincón sample (Fig 4). Chlamydiae was the most abundant phylum sampled for the Mayagüez sample (187 total reads), while Bacteroidetes was the most abundant phylum sampled for the Rincón sample (15 total reads) (Fig 3).

**Fig 4.**
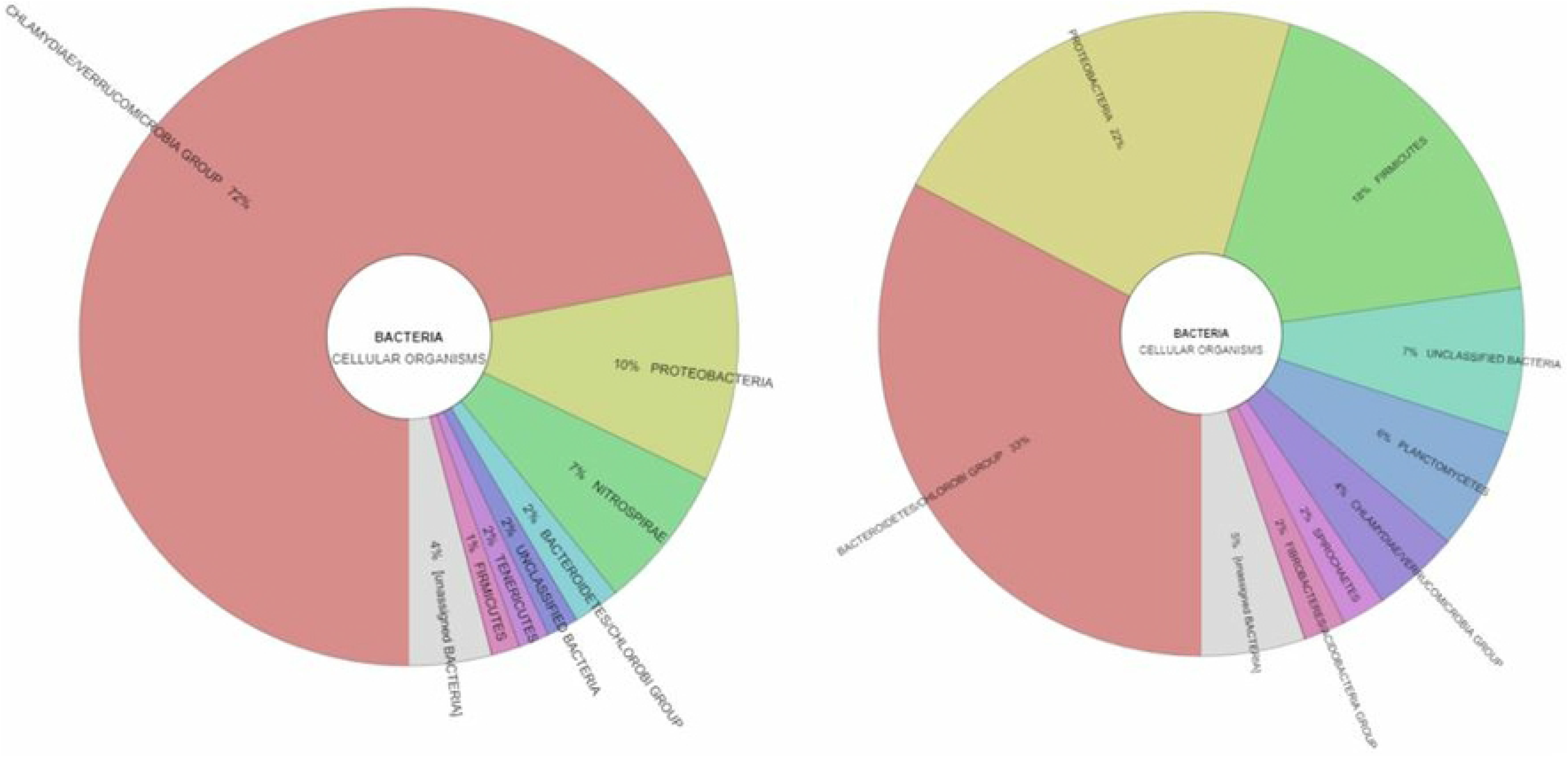
Phylosift results of Bacterial diversity chart from the Mayagüez sample (left) and the Rincón sample (right).

**Fig 5.**
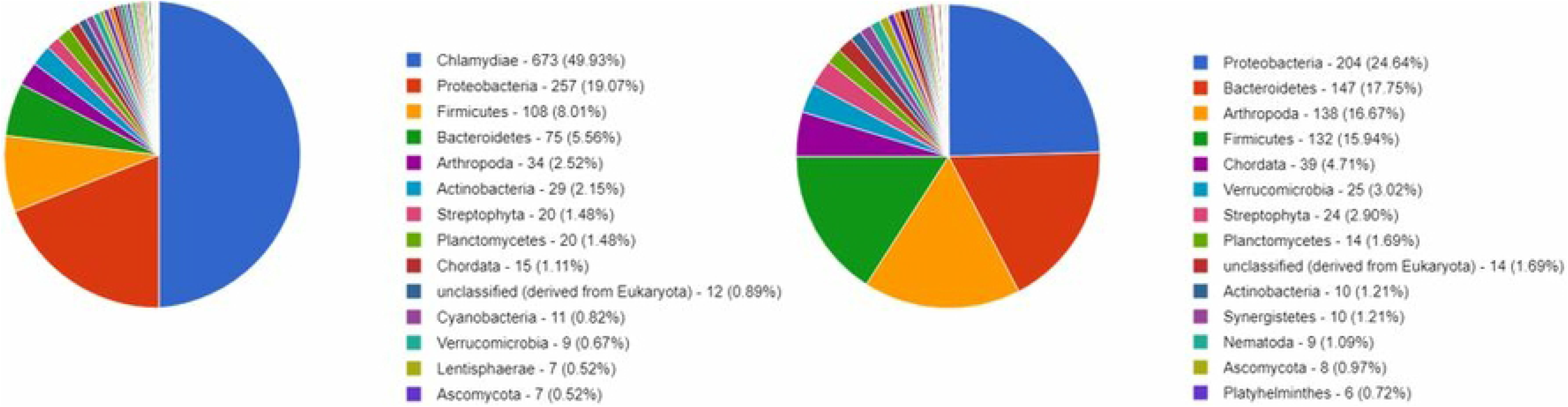
Phylum representation chart for the Mayagüez sample (left) and the Rincón sample (right), created by uploading the sample data to MG-RAST.

**Table 2.** Comparison of annotated reads from the mayor bacterial phyla sampled from *A. monilicornis* guts, based on the program used.

|  | Mayagüez sample |  |  | Rincón sample |  |  |
| --- | --- | --- | --- | --- | --- | --- |
| Phylum<br>(Bacteria) | Phylosift | MEGAN6 | MG-RAST | Phylosift | MEGAN6 | MG-RAST |
| Chlamydia | 187 | 356 | 673 | 2 | 0 | 0 |
| Proteobacteria | 27 | 203 | 257 | 10 | 188 | 204 |
| Bacteroidetes | 6 | 28 | 75 | 15 | 18 | 147 |
| Firmicutes | 4 | 28 | 108 | 8 | 14 | 132 |
| Verrucomicrobia | 0 | 4 | 9 | 0 | 26 | 25 |
| Actinobacteria | 0 | 42 | 29 | 0 | 19 | 10 |
| Planctomycetes | 0 | 2 | 20 | 3 | 2 | 14 |
| Nitrospirae | 18 | 0 | 0 | 0 | 0 | 0 |

**Table 3.** Comparison of annotated reads from the mayor eukaryote phyla sampled from *A. monilicornis* guts, based on the program used.

|  | Mayagüez sample |  |  | Rincón sample |  |  |
| --- | --- | --- | --- | --- | --- | --- |
| **Phylum**<br>**(Eukaryota)** | **Phylosift** | **MEGAN6** | **MG-RAST** | **Phylosift** | **MEGAN6** | **MG-RAST** |
| Arthropoda | 0 | 118 | 34 | 0 | 450 | 138 |
| Chordata | 0 | 4 | 15 | 0 | 12 | 29 |
| Nematoda | 0 | 3 | 0 | 0 | 4 | 9 |
| Streptophyta | 0 | 2 | 20 | 0 | 2 | 24 |
| Ascomycota | 0 | 2 | 7 | 0 | 3 | 8 |
| Alveolata | 24 | 0 | 0 | 1 | 0 | 0 |
| Stramenopiles | 8 | 0 | 0 | 1 | 0 | 0 |

### Eukaryotic reads

According to Phylosift, the two samples had roughly the same number of reads for the protist phyla Alveolata and Stramenopiles (Fig 6). The MG-RAST analysis showed a majority of eukaryotic reads represented by Arthropoda, Streptophyta, and Chordata; the Mayagüez sample had 34 reads for Arthropoda, 20 for Streptophyta, and 15 for Chordata; the Rincón sample had 138 reads for Arthropoda, 39 for Chordata, and 24 for Streptophyta. Finally, MEGAN6 showed the majority eukaryotic reads belonged to Arthropoda, with 118 reads for the Mayaguez sample and 450 reads for the Rincón sample (Table 3).

**Fig 6.**
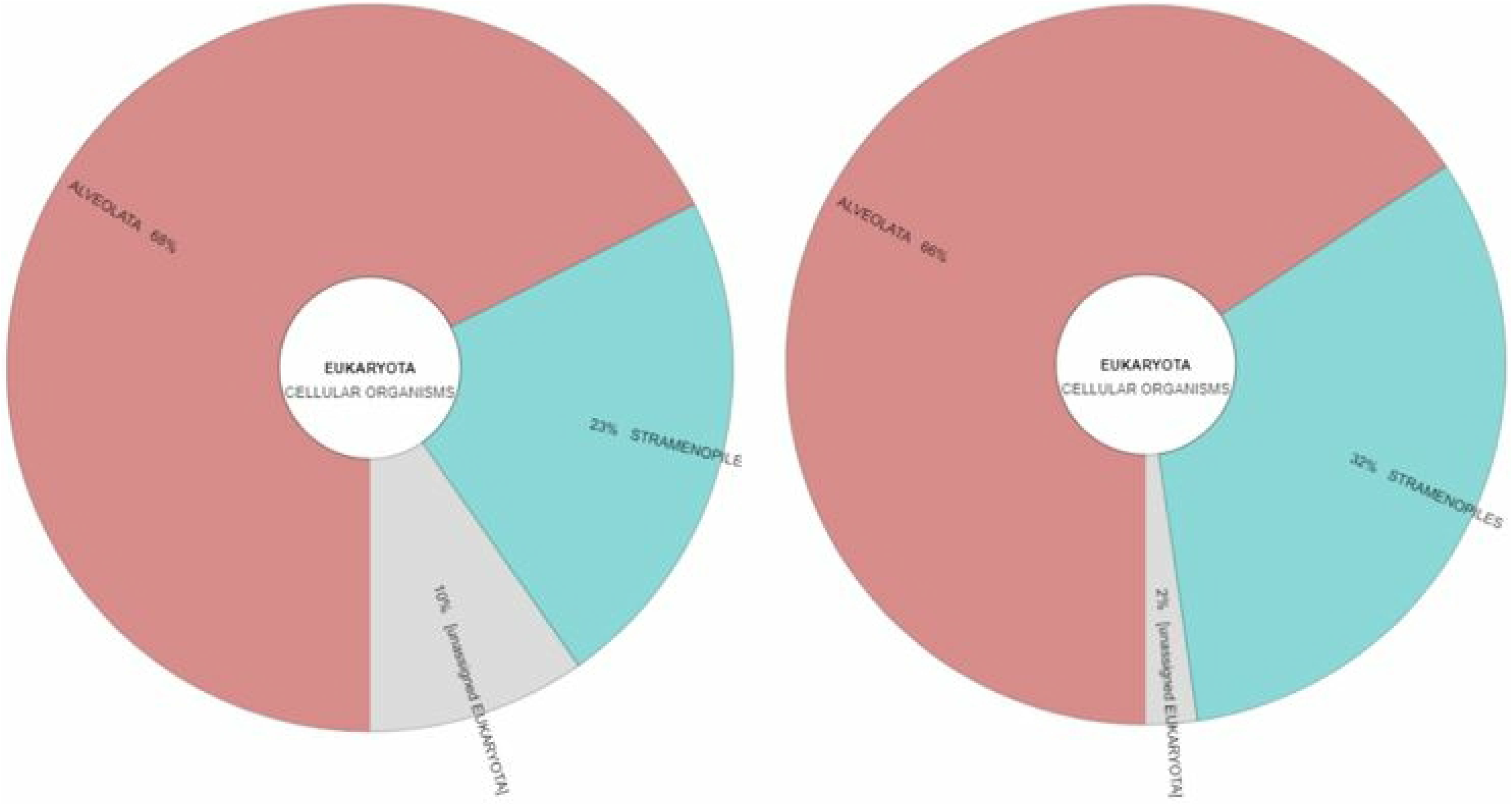
Eukaryotic diversity chart for the Mayagüez sample (left) and the Rincón sample (right), created using Phylosift data. Eukaryotes represented 12% of the sampled reads in the Mayagüez sample, and 4% in the Rincón sample.

### Metabolism annotation

MG-RAST analysis showed that most of the annotated metabolic reads belonged to core cellular metabolism, followed by genetic and environmental metabolic pathways (Fig 7). The metabolic pathways, created via iPath3, can be seen in detail in Figures 8 and 9.

**Fig 7.**
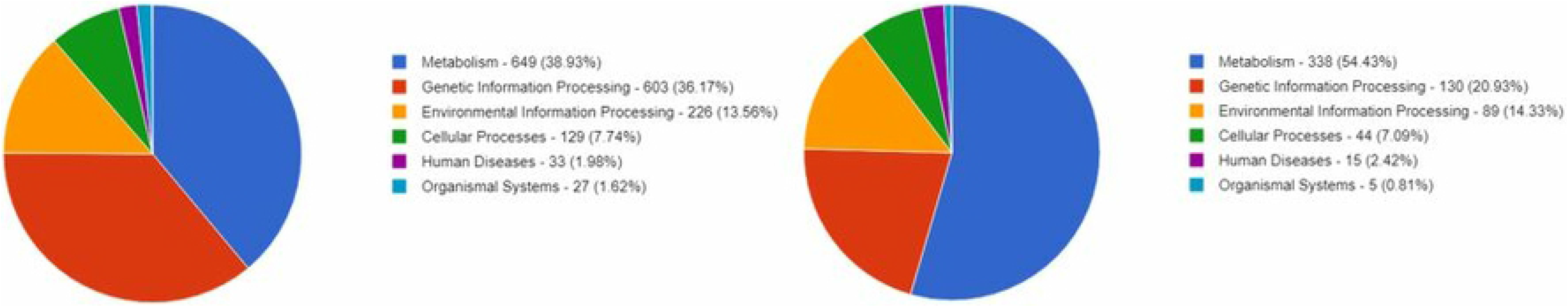
KO metabolism analysis for the Mayagüez sample (left) and the Rincón sample (right), obtained MG-RAST.

**Fig 8.**
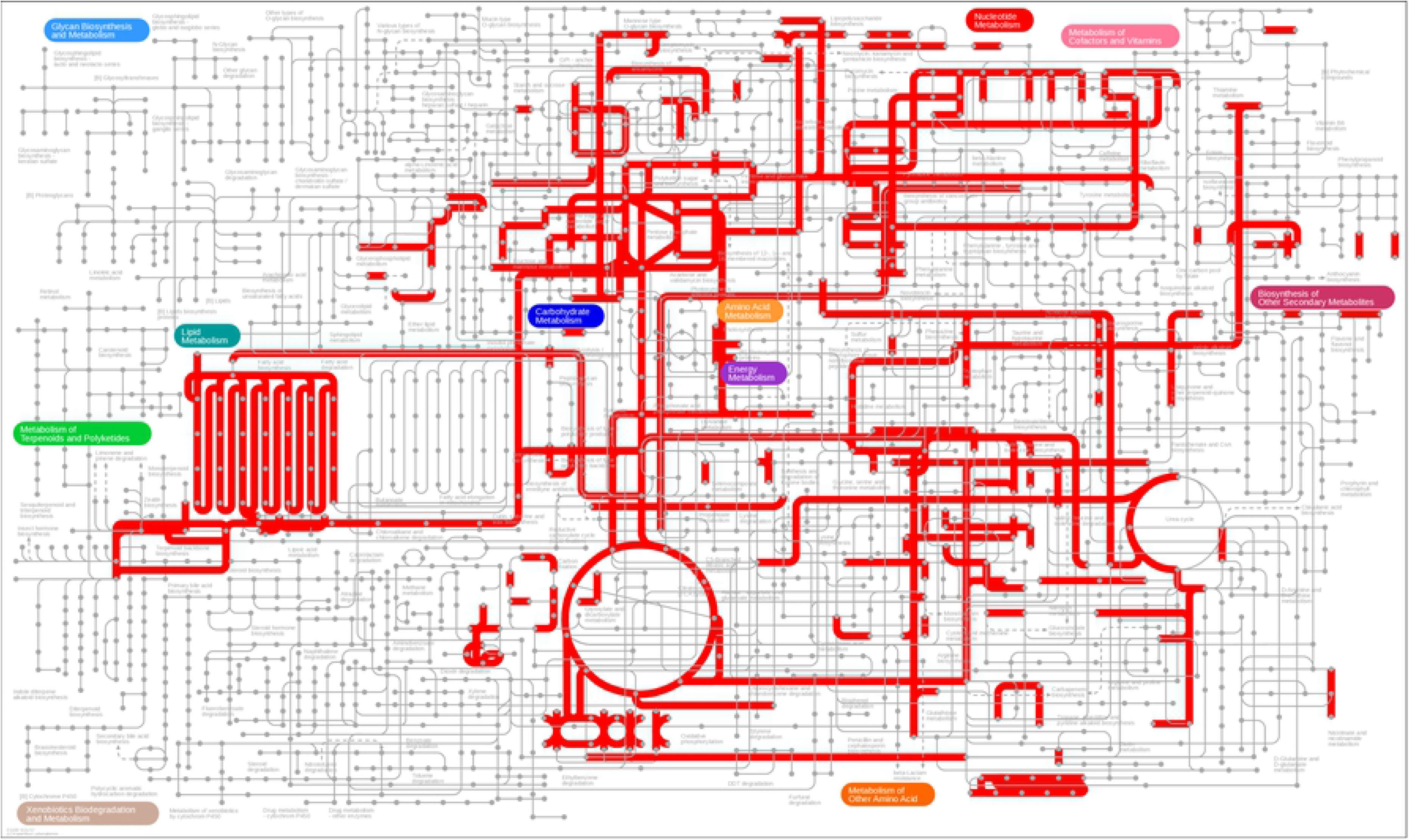
Metabolic pathways for the Mayagüez sample, made by uploading a KO list to iPath3.

**Fig 9.**
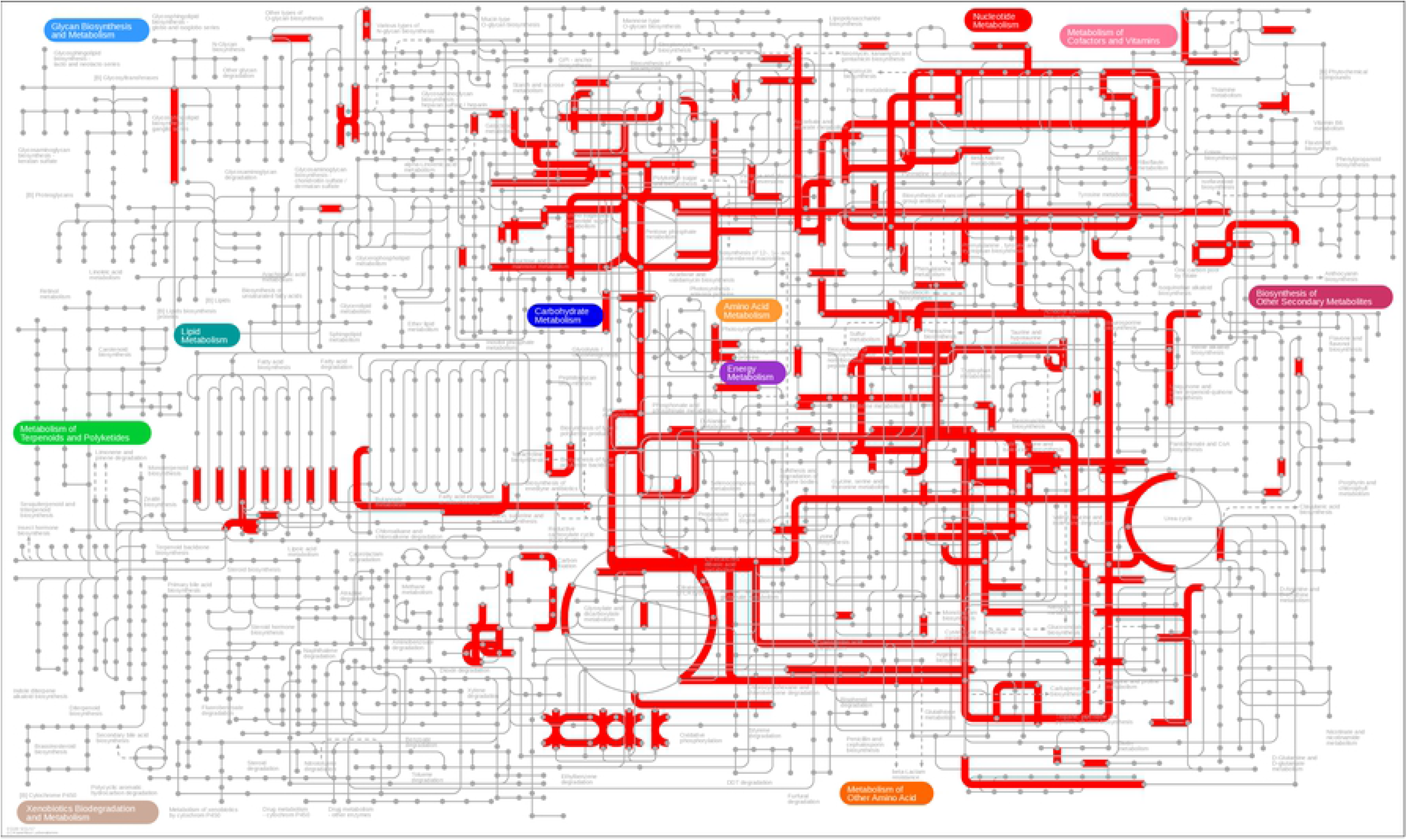
Metabolic pathways for the Rincón sample, made by uploading a KO list to iPath3.

## Discussion

Our results show a fingerprint of the microbial diversity within the gut of *A. monilicornis*. Proteobacteria, Bacteroidetes and Firmicutes were some of the most well represented bacterial phyla in both of our samples (**Table 2**). These findings are comparable with the microbiota found in the guts of termites, which harbor a varying representation of the same phyla as in Rossmassler et al. [9]. Other phyla in common include Actinobacteria and Planctomycetes, hinting at a similar microbial composition between termite and millipede guts [9]. Dittmer et al. [10] found Proteobacteria as one of the most abundant phyla in the terrestrial isopod species they studied. This is to be expected, since both termites and isopods share a similar ecological role to millipedes in the form of terrestrial nutrient recycling, similar to the most abundant phyla found in the present study. Interestingly, it was found that the representation of bacterial phyla in the termites was in part dependent on the gut section sampled; for example, the P1 hindgut compartment was dominated by Firmicutes in most of the termite species studied [9]. Nardi et al. [43] studied the millipede *Cylindroiulus caeruleocinctus* and found that the highest microbial density could be found in the hindgut. Since we sampled the entire gut of the *A. monilicornis* specimens, we cannot determine any differences in representation across the millipede’s gut sections. However, it could be an interesting topic for future studies.

Gut microbial diversity could potentially vary between different populations of millipedes. Dittmer et al. [10] examined the terrestrial isopod *Armadillidium vulgare* finding differences in microbial communities based on localities. Our data may echo those results, since the prevailing bacterial phyla between our two samples differed (**Table 2**). This could have been due to just undersampling in our study but, these differences could also be due to changes in diet; the termite study as in Rossmassler et al. [9]. For example, that the phylum Bacteroidetes was most abundant in wood-feeding termites [9]. In contrast, Knapp et al. [15] who studied the effect of different diets on the gut microbiota of the alpine millipede *Cylindroiulus fulviceps*, found no significant microbial diversity changes between the samples. In our case, we added an additional variable to our experiment in the form of starvation time of the millipedes. The Mayagüez millipede sample was starved for approximately 24 hours; the Rincón millipede sample was starved for ten days. Due to the small sample size studied and variable conditions prior to gut extraction, some questions remain to be answered by the present study: Did starving cause differences in gut microbial diversity? Or was it differences in soil microbial composition between the two sites? Or a combination of both? A greater sample size could have allowed us to answer these questions with some degree of statistical significance.

Several additional questions arise from this study. Dittmer et al. [10] found that 70% of the gut microbial taxa found in the isopod *Armadillidium vulgare* were also detected in feces and in the soil, suggesting that an important fraction of the microbiota may be acquired from environmental sources. The question of how millipedes acquire their microbiota, whether from the soil they inhabit, or maternal transmission from other millipedes, or a combination of both still remain largely unanswered. Crawford et al. [2] found that the lumen bacteria of the desert millipede *Orthoporus ornatus* virtually disappeared after molting. Does microbial diversity representation change across a millipede’s lifespan, before and after molting? These questions, however, are beyond the scope of this study and require broader geographic sampling.

It is very odd that we did not get many annotated Nematoda reads, for example with Phylosift we were unable to obtain any Nematoda reads (**Table 3**). We expected to find more, since we visually identified nematodes inside the extracted guts before sequencing them. We also expected a larger number of reads for the fungal phylum Ascomycota; according to the work of Byzov et al. [44], ascomycetes are the most common yeast strains found on certain species of millipedes. However, we obtained a small number of Ascomycota reads compared to other phyla (**Table 3**). With our current data, we are not able to determine why this happened again this is probably due to under sampling.

Some of the reads seem to be misannotated as well. For example, eukaryotic reads analyzed by Phylosift were linked to the protists Alveolata and Stramenopiles; specifically, the aquatic species *Thalassiosira sp., Kryptoperidinium foliaceum* and *Durinskia baltica*. This could be interpreted as Phylosift miss-annotating the eukaryotic reads to the organisms available in the database with the closest genetic resemblance. *Thalassiosira*, for example, is notable for being one of the first marine phytoplankton genera whose genome was sequenced [45]; perhaps most of the protist reads databased belong to a small number of genera or species. On the other hand, the MG-RAST and MEGAN6 analyses annotated some of the eukaryote reads in both samples as belonging to Chordata (Fig 2, Fig 5), and MG-RAST annotation for metabolism returned reads related to human diseases (Fig 7). These could have been miss-annotations, or some presence of contaminants in the samples during DNA extraction. Errors in annotation have been a problem since the advent of sequencing, and according to Schnoes et al. [46] it has “increased from 1993 to 2005” for public databases in particular. Indeed, studies have reported error rates as high as 90% for protein and rRNA sequences, for example, in databases such as GenBank and TrEMBL [45–46]. Though this topic is beyond the scope of this study, we believe finding techniques to consistently circumvent these annotation errors is of upmost importance to the omics fields.

It is worth noting that most arthropod metagenomic studies have been done on insects [28]. Focusing on lesser studied arthropods, millipedes included, could help broaden our understanding of host microbiota diversity and coevolution. With regards to the present study, a greater sample size could have allowed us to answer more questions, it nevertheless showed what to expect from sequencing the gut of a millipede using nanopore sequencing. We hope to be able to continue studying millipede gut metagenomics or to encourage more studies from the community of metagenomics researchers to continue further work. Future directions could focus on a more standardized sampling and extraction protocol, to be able to secure a larger sample size, and consequently compile a greater amount of the microbial taxa found in the gut. With more samples and sequencing depth, we could better determine any differences in microbial diversity between populations across Puerto Rico and delve deeper into the ecological importance of millipedes in the biomes they inhabit.

## Acknowledgements

Authors thank PSC staff Tom Maiden, Rick Costa, and Roberto Gomez for help with installing albacore on the Bridges system. Laboratory work was funded by a NIH PR-INBRE Grant Contract #5P20GM102475 sub-award to AVD. Bioinformatics was funded by an NSF-XSEDE Grant TG-BIO170059 awarded to AVD. Publication costs were supported by NIFA HSI Grant (2018-38422-28612) awarded to AVD.

Author Contributions
AVD conceived the project; AVD, MJC, AR and CJSF contributed resources; AVD and OJGC analyzed the data; OJGC, AVD, MJC, and CJSF wrote the paper.

## Literature Cited

1. Taylor E (1982) Role of Aerobic Microbial Populations in Cellulose Digestion by Desert Millipedes. Appl Environ Microbiol 44: 281–291.

2. Crawford C, Minion G, Boyers M (1983) Intima morphology, bacterial morphotypes, and effects of annual molt on microflora in the hindgut of the desert millipede, Orthoporus ornatus (Girard) (Diplopoda:Spirostreptidae). Int J Insect Morphol Embriol 12: 301–312.

3. Vélez M (2014) Los Gongolies, Gungulenes o Milpiés (Clase Diplopoda). In Joglar, R., Santos-Flores, C. & Torres-Pérez, J. Biodiversidad de Puerto Rico: Invertebrados. pp. 240–253.

4. Snyder B, Hendrix P (2008) Current and Potential Roles of Soil Macroinvertebrates (Earthworms, Millipedes, and Isopods) in Ecological Restoration. Restor Ecol 16: 629–636.

5. Pitz K, Sierwald P (2010) Phylogeny of the millipede order Spirobolida (Arthropoda: Diplopoda: Helminthomorpha). Cladistics 26: 497–525.

6. Kitz F, Steinwandter M, Traugott M, Seeber J (2015) Increased decomposer diversity accelerates and potentially stabilises litter decomposition. Soil Biol Biochem 83: 138–141. doi:10.1016/j.soilbio.2015.01.026.

7. Sierwald P, Bond JE (2007) Current Status of the Myriapod Class Diplopoda (Millipedes): Taxonomic Diversity and Phylogeny. Annu Rev Entomol 52: 401–420. Available: http://www.annualreviews.org/doi/10.1146/annurev.ento.52.111805.090210. Accessed 28 February 2018.

8. Brewer MS, Sierwald P, Bond JE (2012) Millipede Taxonomy after 250 Years: Classification and Taxonomic Practices in a Mega-Diverse yet Understudied Arthropod Group. PLoS One 7: e37240.

9. Rossmassler K, Dietrich C, Thompson C, Mikaelyan A, Nonoh J, et al. (2015) Metagenomic analysis of the microbiota in the highly compartmented hindguts of six wood- or soil-feeding higher termites. Microbiome 3: 111–118.

10. Dittmer J, Lesobre J, Moumen B, Bouchon D (2016) Host origin and tissue microhabitat shaping the microbiota of the terrestrial isopod Armadillidium vulgare. FEMS Microbiol Ecol 92: fiw063. doi:10.1093/femsec/fiw063.

11. Szabo I, Nasser E, Striganova B, Rakhmo Y, Jager K, et al. (1990) Interactions among Millipedes (Diplopoda) and their Gut Bacteria. Ber nat-med Verein Innsbruck 10: 289–296.

12. Byzov B (2006) Gut Microbiota of Millipedes. Konig & Varma, A. Gut Microorganisms of Termites and Other Invertebrates. Heidelberg: Springer. pp. 89–114.

13. Konig H (2006) Bacillus species in the intestine of termites and other soil invertebrates. J Appl Microbiol 101: 620–627.

14. Sustr V, Chronakova A, Semanova S, Tajovsky K, Simek M (2014) Methane Production and Methanogenic Archaea in the Digestive Tracts of Millipedes (Diplopoda). PLoS One 9: 7. doi:10.1371/journal.pone.0102659.

15. Knapp B, Seeber J, Podmirseg S, Rief A, Meyer E, et al. (2009) Molecular fingerprinting analysis of gut microbiota of Cylindroiulus fulviceps (Diplopoda). Pedobiologia (Jena) 52: 325–336.

16. David J (2014) The role of litter-feeding macroarthropods in decomposition processes: a reappraisal of common views. Soil Biol Biochem 76: 109–118.

17. Gabel K, Hunsberger A, Mannion C, Buss L, Buss E (2006) Yellow-banded Millipede. Gainesville. Available: http://trec.ifas.ufl.edu/mannion/pdfs/Yellow-bandedMillipede.pdf.

18. Shelley R (2014) A consolidated account of the polymorphic Caribbean milliped, Anadenobolus monilicornis (Porat, 1876) (Spirobolida: Rhinocricidae), with illustrations of the holotype. Insecta mundi 0378: 1–12.

19. Contreras K, Cafaro M (2013) Morphometric Studies in Enterobryus luteovirgatus sp. nov. (Ichthyosporea: Eccrinales) Associated with Yellowbanded Millipedes in Puerto Rico. Acta Protozool 52: 291–297.

20. Thomas T, Gilbert J, Meyer F (2012) Metagenomics – a guide from sampling data analysis. Microb Inform Exp 2: 3.

21. Qi W, Nong G, Preston JF, Ben-Ami F, Ebert D (2009) Comparative metagenomics of Daphnia symbionts. BMC Genomics. doi:10.1109/TCSI.2002.800838.

22. Robidart JC (2006) Metagenomics of the Riftia pachyptila symbiont University of California, San Diego.

23. Warnecke F, Luginbühl P, Ivanova N, Ghassemian M, Richardson TH, et al. (2007) Metagenomic and functional analysis of hindgut microbiota of a wood-feeding higher termite. Nature 450: 560–565. doi:10.1038/nature06269.

24. Engel P, Martinson V, Moran N (2012) Functional diversity within the simple gut microbiota of the honey bee. PNAS 109: 11002–11007.

25. Yun J, Roh S, Whon T, Jung M, Kim M, et al. (2014) Insect Gut Bacterial Diversity Determined by Environmental Habitat, Diet, Developmental Stage, and Phylogeny of Host. Appl Environ Microbiol 80: 5254–5264.

26. Chen B, Teh B, Sun C, Hu S, Lu X, et al. (2016) Biodiversity and Activity of the Gut Microbiota across the Life History of the Insect Herbivore Spodoptera littoralis. Sci Rep 6: 29505. doi:10.1038/srep29505.

27. Muturi E, Ramirez J, Rooney A, Kim C (2017) Comparative analysis of gut microbiota of mosquito communities in central Illinois. PLoS Negl Trop Dis 11. doi:10.1371/.

28. Bouchon D, Zimmer M, Dittmer J (2016) The terrestrial isopod microbiome: An all-in-one toolbox for animal-microbe interactions of ecological relevance. Front Microbiol 7: 1–19. doi:10.3389/fmicb.2016.01472.

29. Degli M, Martinez E (2017) The functional microbiome of arthropods. PLoS One 12. doi:https://doi.org/10.1371/journal.pone.0176573.

30. Jain M, Olsen HE, Paten B, Akeson M (2016) The Oxford Nanopore MinION: Delivery of nanopore sequencing to the genomics community. Genome Biol 17: 1–11. doi:10.1186/s13059-016-1103-0.

31. Technologies ON (2017) Albacore basecaller from Oxford Nanopore. Available: https://mirror.oxfordnanoportal.com/software/analysis/ont_albacore.

32. Technologies ON (2017) New basecaller now performs ‘raw basecalling’, for improved sequencing accuracy. Available: https://nanoporetech.com/about-us/news/new-basecaller-now-performs-raw-basecalling-improved-sequencing-accuracy. Accessed 20 September 2017.

33. Schultz D n. d. (2018) Pauvre: QC and genome browser plotting Oxford Nanopore and PacBio long reads. Available: https://github.com/conchoecia/pauvre.

34. Bolger AM, Lohse M, Usadel B (2014) Trimmomatic: a flexible trimmer for Illumina sequence data. Bioinformatics 30: 2114–2120. Available: https://academic.oup.com/bioinformatics/article-lookup/doi/10.1093/bioinformatics/btu170. Accessed 14 September 2017.

35. Chikhi R, Medvedev P (2014) Informed and automated k-mer size selection for genome assembly. Bioinformatics 20: 31–37.

36. Zerbino DR, Birney E (2008) Velvet: algorithms for de novo short read assembly using de Bruijn graphs. Genome Res 18: 821–829. Available: http://www.genome.org/cgi/doi/10.1101/gr.074492.107. Accessed 14 September 2017.

37. Koren S, Walenz B, Berlin K, Miller J, Bergman N, et al. (2017) Canu: scalable and accurate long-read assembly via adaptive k-mer weighting and repeat separation. Genome Res 27: 722–736.

38. Darling AE, Jospin G, Lowe E, Matsen FA, Bik HM, et al. (2014) PhyloSift: phylogenetic analysis of genomes and metagenomes. PeerJ 2: e243. Available: https://peerj.com/articles/243. Accessed 1 March 2018.

39. Altschul S, Gish W, Miller W, Myers E, Lipman D (1990) Basic local alignment search tool. J Mol Biol 215: 403–410.

40. Huson D, Auch A, Qi J, Schuster S (2007) MEGAN analysis of metagenomic data. Genome Res 17: 377–386.

41. Meyer F, Paarmann D, D’Souza M, Olson R, Glass E, et al. (2008) The metagenomics RAST server - a public resource for the automatic phylogenetic and functional analysis of metagenomes. BMC Informatics 9. doi:10.1186/1471-2105-9-386.

42. Yamada T, Letunic I, Okuda S, Kanehisa M, Bork P (2011) iPath2.0: interactive pathway explorer. Nucleic Acids Res 39: 412–415.

43. Nardi J, Bee C, Taylor S (2016) Compartmentalization of microbial communities that inhabit the hindguts of millipedes. Arthropod Struct Dev 45: 462–474.

44. Byzov B, Thanh V, Babjeva I (1993) Yeasts associated with soil invertebrates. Biol Fertil Soils 16: 183–187.

45. Armbrust E, Berges J, Bowler C, Green B, Martinez D, et al. (2004) The Genome of the Diatom Thalassiosira Pseudonana: Ecology, Evolution, and Metabolism. Science (80-) 306: 79–86. doi:10.1126/science.1101156.

46. Schnoes AM, Brown SD, Dodevski I, Babbitt PC (2009) Annotation error in public databases: Misannotation of molecular function in enzyme superfamilies. PLoS Comput Biol 5. doi:10.1371/journal.pcbi.1000605.

47. Tripp HJ, Hewson I, Boyarsky S, Stuart JM, Zehr JP (2011) Misannotations of rRNA can now generate 90 false positive protein matches in metatranscriptomic studies. Nucleic Acids Res 39: 8792–8802. doi:10.1093/nar/gkr576.

